# Evaluating DCA-based method performances for RNA contact prediction by a well-curated dataset

**DOI:** 10.1101/822023

**Authors:** F. Pucci, M. Zerihun, E. Peter, A. Schug

**Affiliations:** John von Neumann Institute for Computing, Jülich Supercomputer Centre, Forschungszentrum Jülich, 52428 Jülich, Germany; Steinbuch Centre for Computing, Karlsruhe Institute of Technology, 76344 Eggenstein-Leopoldshafen, Germany; Department of Physics, Karlsruhe Institute of Technology, 76344 Eggenstein-Leopoldshafen, Germany

**Author notes:** These authors contribute equally to this work (F. Pucci, M. Zerihun).

## Abstract

RNA molecules play many pivotal roles in the cellular functioning that are still not fully understood. Any detailed understanding of RNA function requires knowledge of its three-dimensional structure, yet experimental RNA structure resolution remains demanding. Recent advances in sequencing provide unprecedented amounts of sequence data that can be statistically analysed by methods such as Direct Coupling Analysis (DCA) to determine spatial proximity or contacts of specific nucleic acid pairs, which improve the quality of structure prediction. To quantify this structure prediction improvement, we here present a well curated dataset of about seventy RNA structures with high resolution and compare different nucleotide-nucleotide contact prediction methods available in the literature. We observe only minor difference between the performances of the different methods. Moreover, we discuss how these predictions are robust for different contact definitions and how strongly depend on procedures used to curate and align the families of homologous RNA sequences.

## Introduction

RNA molecules play fundamental roles in a large variety of processes within cells. For example messenger RNAs (mRNAs) carry the genetic information akin to blueprint for protein synthesis, transfer RNA (tRNA) then carry specific amino acids during protein synthesis to the site of protein elongation[1]. More recently other tasks of RNA were identified, such as noncoding RNAs (ncRNAs) fulfilling fundamental roles in the control of gene expression [2, 3] or small interference RNAs (siRNAs) and micro RNAs (miRNAs) that can regulate and repress the expression of target gene by interfering with the transcriptional regulation [4, 5].

Long non-coding RNAs (lncRNAs) also contribute to these modulation mechanisms even if they are less understood. Metabolite-binding RNA structures called riboswitches that belong to the 5’ untranslated regions (5’-UTR) of the mRNA bind selectively and with high affinity small molecules and this biding induces major conformation rearrangements of the three-dimensional structure of the riboswitches. The two competing conformations can inhibit or activate the expression of the target gene by interfering with the translation regulation. The study of lncRNAs is of particular high interest as they are frequently involved in pathogenic mechanisms and thus can be targeted for therapeutic strategies [6].

To truly understand the molecular mechanisms of lncRNAs and their function, it is important to know their three-dimensional structure. Experimental techniques to determine the 3d structure include “classical” methods such as X-ray diffraction crystallography or nuclear magnetic resonance (NMR), which provide direct structural information. Other methods do not directly provide structural information but have first to be carefully interpreted (e.g. small-angle scattering (SAXS)[7] or Fluorescence Resonance Energy Transfer (FRET) [8]). Yet in spite of considerable progress of experimental techniques, the number of structurally resolved RNA structures collected in public databases [9, 10] is still small due to experimental limitations and considerably lags the number of known sequences.

Computational methods contributed substantially to decipher how RNA structure and dynamics determine its functions [11, 12, 13]. A series of computational tools have been developed to predict the RNA structure from the sequence using different approaches that can be roughly divided in fragment-based, physics-based and comparative modeling [14, 15, 16, 17, 18, 19, 20, 21, 22, 23, 24, 25, 26, 27]. Their performances are improving as one can see from the results of the three RNA-puzzle rounds [28, 29, 30] where a set of experimentally resolved 3D structures has been blindly predicted.

Recent investigations [26, 31, 32] have shown that the performances of these methods can be substantially improved by using information extracted from multiple sequence alignment (MSA) of families of homologous RNAs. Improvements are achieved by identifying top-ranked site-pairs with stronger coevolutionary signals and using them as distance constraints in modeling tools.

Thanks to the advancement of next generation sequencing technologies, the huge and increasing amount of sequence data available can be fully exploited to study and model RNA structures.

To get more insights in these issues, in this paper we set up a manually curated dataset of about seventy RNA structures with high resolution and evaluate the performances of different contact prediction methods on this set. Moreover, we analyze the impact of important features on their performances such as the effective number of homologous RNA sequence that are available, the nucleotide-nucleotide contact definition and the procedure to construct, align and curate the MSA.

## Methods

### Dataset curation

We manually curated a dataset of three-dimensional RNA structures starting our analysis from the whole Protein Data Bank [9] and selecting all RNA structures that satisfied the following criteria :

- the RNA structures were not in a complex with proteins or DNAs
- only monomeric structures were considered
- the length of the RNA sequences had to be bigger than 40 nucleic bases
- in cases of structures with similar sequences (sequence identity (SI) between pairs of sequences of 50%) we choose the structures with higher resolution
- only structures resolved via X-ray crystallography were taken into account with resolution below 3.6*Å*

We associated one RNA family from the Rfam database [33] to each entry in the dataset by choosing the family with the highest match to the sequence. This search has been done employing INFERence of RNA ALignment tool (Infernal) using the BIT value as a match score [34].

For each family we then computed the number of effective sequences *M*_*eff*_ via the pydca software package [35]. This value is computed from the alignment of the given family as *M*_*eff*_ = ∑_*k*_ *ω*_*k*_ where *ω*_*k*_ is the weight of the *k*^*th*^ entry in the given cluster of similar sequences that is identified using a cut-off on the SI equal to 0.8 [36].

We further split the final set *𝒟* comprised by 69 RNA structures into two subsets : *𝒟*^*High*^ containing 36 structures associated to RNA families with a *M*_*eff*_ larger than 70. The 33 remaining entries belong to *𝒟*^*Low*^set and have a MSA with *M*_*eff*_ *<* 70. The list of all entries in *𝒟* with their characteristics is reported in the Table 1 of the Supplementary information.

**Table 1:** Positive predicted values (PPV) according to the type contact considered.

|  | A | C | G | U |
| --- | --- | --- | --- | --- |
| A | 18% | 10% | 21% | 64% |
| C | 10% | 6% | 75% | 3% |
| G | 21% | 75% | 17% | 32% |
| U | 64% | 3% | 32% | 10% |

### Contact definition

In order to study how the nucleotide-nucleotide contact definition influences the performance of DCA methods, we computed and compared positive predicted values (PPVs) using different criteria to construct the contact maps from PDB structures. We choose and tested six distance-based criteria to classify if a pair of nucleotides is in direct physical interaction or not:

1. Two nucleotides are in contact if the distance between the N9 atoms of a purine or the N1 of a pyrimidine is smaller than 9.5 Å.
2. Two nucleotides are in contact if the distance between their C1’ atoms is smaller than 12.0 Å.
3. Two nucleotides are in contact if the distance between two of any of their heavy atoms is smaller than 3.5, 5.5, 7.5 and 9.5 Å.

### Multiple sequence alignment and curation

As the quality of the multiple sequence alignment critically impacts the accuracy of the subsequent contact prediction, we tested different methods to perform and curate such alignments. We then studied how the mean-field DCA performances changed according to the method used.

1. **Search**. We started to verify if the construction of the RFAM families can influence the prediction performance. To do that we used either the RFAM families as given in RFAM v14.1, but we also reconstruct them using both less and more stringent E-value cut-offs equal to 0.0001 and 0.99 respectively. This search is done using the Infernal software (cmsearch) and the precomputed covariance model (CM) of the given family.
2. **Align**. The first methods used to perform the MSA of the RFAM families is the Infernal software [34]. More in details all entries of the RFAM family considered are aligned using the corresponding covariance model (CM) that is a specific profile stochastic context-free grammar that scores a combination of sequences and RNA secondary structure consensus. We tested also other three commonly used tools for the multiple sequence alignment of RNAs that are CLUSTALW [37], MUSCLE [38] and MAAFT [39].
3. **Trim**. From the MSA of the given family we test three different possibilities : in the first one only positions corresponding to the target sequence were considered for the DCA computations; in the second and the third ones, before the DCA computation we trimmed the MSA by selecting only columns that have less than 50% and 20% of gaps respectively.

### Coevolution-based methods

Different methods have been developed for the implementation of the direct coupling analysis (DCA) of RNAs. Given a family of homologous RNA sequences, these statistical models assigns probabilities *P* (*S*) to each sequence *S* = *a*_1_*a*_2_…*a*_*L*_ of length *L* using the the Boltzmann law as

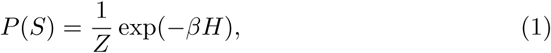

where *β* is the inverse of the temperature usually fixed to one without loss of generality, *Z* the partition function and *H* the Hamiltonian taken of the form

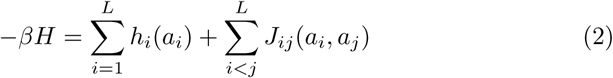

that contains single site terms, *i*.*e. h*_*i*_(*a*_*i*_), and nucleotide pair interactions *J*_*ij*_(*a*_*i*_, *a*_*j*_). In DCA these parameters are inferred from the input MSA using different approaches that are briefly shown here. See also the references [40, 41] for recent reviews on the topic.

- **Mean-field DCA**. Here, a standard mean-field approximation of the partition function is done in order to obtained the couplings and the singlesite fields in a computational efficient way. Within this approximation the *J*_*ij*_(*a*_*i*_, *a*_*j*_) are obtained as

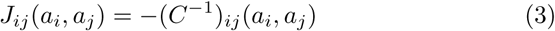

where *C* is the matrix of correlations whose elements are given by *C*_*ij*_(*a*_*i*_, *a*_*j*_) = *f*_*ij*_(*a*_*i*_, *a*_*j*_)− *f*_*i*_(*a*_*i*_)*f*_*j*_(*a*_*j*_) with *f*_*ij*_(*a*_*i*_, *a*_*j*_) and *f*_*i*_(*a*_*i*_) the empirical frequency counts obtained from the MSA columns. The single-site fields *h*_*i*_(*a*_*i*_) are obtained self-consistently from the frequencies *f*_*i*_(*a*_*i*_) and the couplings in eq (3). We used the mean-field implementation in pydca [35].
- **Boltzmann Learning**. In this statistical approach the parameter *J*_*ij*_(*a*_*i*_, *a*_*j*_) and *h*_*i*_(*a*_*i*_) are obtained from the minimization of the negative log likelihood

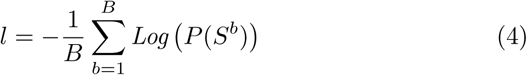

where *P* (*S*^*b*^) (*b* = 1…*B*) is a set of independent equilibrium configurations of the model, i.e. RNA sequences that belong to a MSA. A direct way to solve the problem is to do a “brute-force” minimization starting from an initial guess for the values of the couplings and fields and using a gradient descent algorithm that employs a Markov chain Monte Carlo method for the gradient evaluation. For more details on the method and the implementation that we used see [42].
- We use the **EVcouplings** [31] implementation that exploit a pseudolikelihood maximization direct couplings analysis (plmDCA) [43]. In this method the probability in eq (4) can be substituted with the conditional probability of observing one variable *a*_*r*_ given the observation of the others 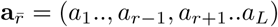. Given the MSA, one has then to minimize the conditional log likelihood

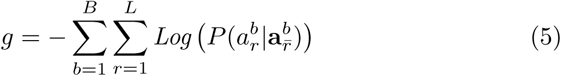

with regularization to estimate the couplings and the fields. This strategy, while retains the accuracy of the full likelihood approach, greatly increases its computational efficiency.
- **GREMLIN** [44]employs a learning procedure that is based on the pseudolikelihood optimization. However GREMLIN can incorporate prior information on predicted secondary structure and on sequence separation. This incorporation allows the method to be likely more accurate when the number of aligned sequences is limited [44].
- **CCMpred** [45] is an implementation based on plmDCA similar to EV-Fold. CCMPred is computationally optimized for GPU architectures.
- **PSICOV**[46] computes the so-called precision matrix defined as

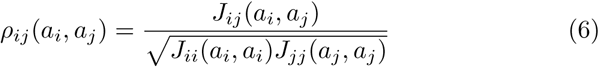

in terms of the inverse of the covariance matrix (3). *ρ*_*ij*_ encodes the correlation between any pair of amino acids or nucleotides at two sites, in terms of the frequencies at all other sites and identify which pairs are likely to be in direct physical contact in the native structure. The estimation of the inverse covariance matrix is done employing a graphical Lasso approach and the final PSICOV score is given by 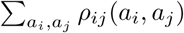 followed by an average product correction (APC) [47].

## Results

### Assessing the performance of DCA-based methods

In this section we compare the performance of the prediction methods tested, namely the mean-field of pydca [35], EVcouplings [31], Boltzmann learning [42], GREMLIN [44], CCMpred [45] and PSICOV [46]. In fig. (1.a) we report the positive predicted values (PPV) on the dataset *𝒟* as a function of the number of contacts. Here we are considering all contacts in the PDB structures that are distant in the sequence more than four nucleotides.

**Figure 1:**
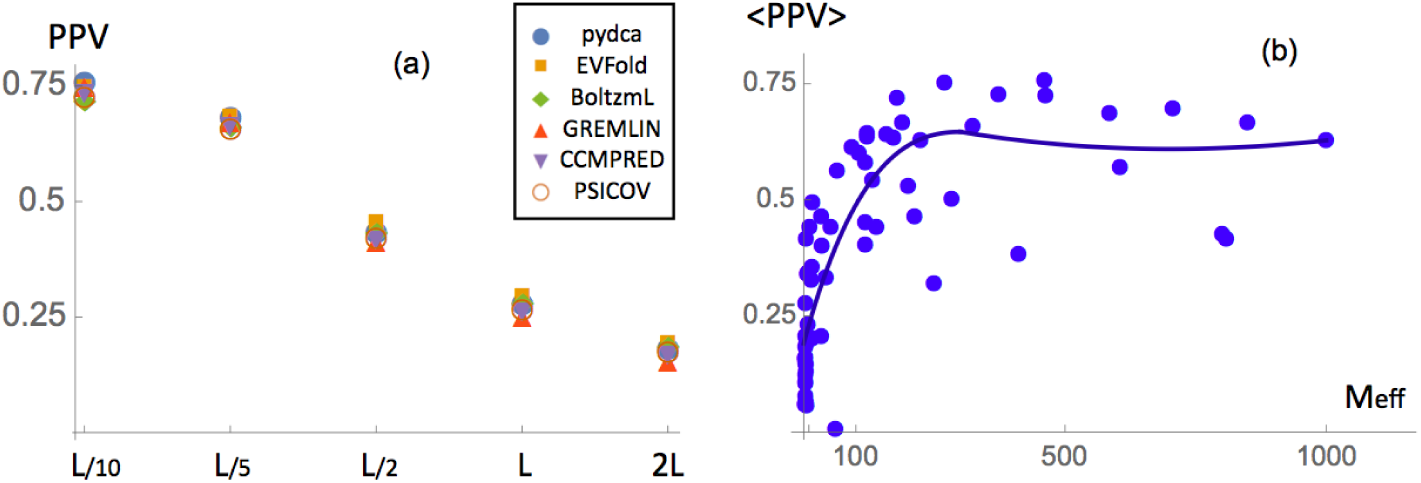
(a) Prediction performances of the different methods analyzed in this papers by PPV as a function of the number of top scoring contacts. All contacts that are separated along the sequence by at least 4 nucleotides are considered. (b) Averaged PPV of all prediction methods as a function of the effective number of sequences *M*_*eff*_.

The performance based on PPV are generally quite good. We find PPV on the order of 75% for the top *L*/10 contacts that goes smoothly down to 0.25% if one considers the top *L* contacts. Among all methods no statistically significant differences can be observed as measured from the Kolmogorov-Smirnov test of the different prediction results. A slightly more accurate performance for small number of contact can be observed for pydca and GREMLIN for *L*/10 number of contacts while at *L*/2 contacts the EVcouplings is few percents more accurate than other predictors (see also Table 1).

When performances are evaluated on *𝒟*^*H*^, *i*.*e*. the set of PDBs associated to RFAM families with *M*_*eff*_ ≥ 70, the performances reach higher PPV equal to about 60% (at *L*/2) in contrast with a PPV of 26% in the *𝒟*^*L*^ set in which only families with *M*_*eff*_ *<* 70 are considered (see fig.2 and Table 1).

**Figure 2:**
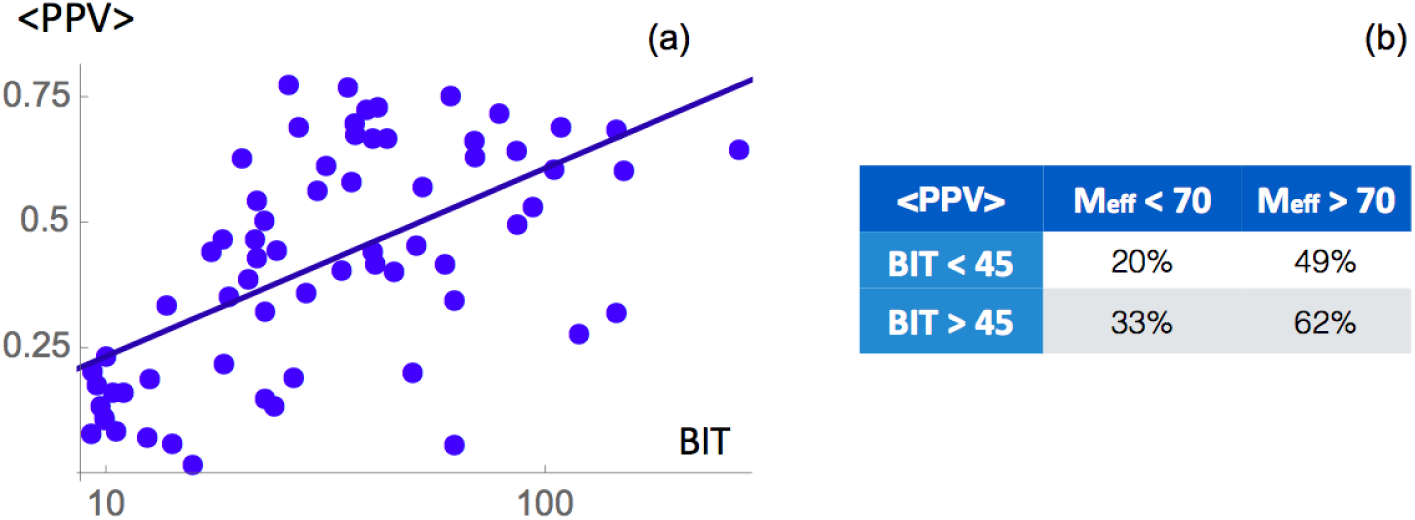
(a) Averaged PPV of all prediction methods as a function of the BIT value for the chosen RFAM family. (b) Table of comparison for PPV as influenced by different *M*_*eff*_ and *BIT*.

In fig. (1.b) we investigate how performances are related to *M*_*eff*_ by plotting the average PPV rate of different methods versus the *M*_*eff*_ of the given RFAM family. We observe a clear growth of the prediction accuracy up to *M*_*eff*_ values equal to about 200 while above that threshold the performances stay approximately constant. This behavior could be due to the fact that families with increasing number of sequences tend to have higher probability of misannotations and biases and thus after that *M*_*eff*_ threshold the additional information and noise are both added to the MSA.

performances are not only related to the *M*_*eff*_ of RFAM families but also from how well the target sequence aligns. In order to check this dependence in fig. (2.a) we plotted the averaged PPV with the BIT value computed from Infernal, a score measuring the probability of the query sequence to match the covariance model. We observe a linear relation between these two quantities. To check the effect of both *M*_*eff*_ and the BIT score, we first divided each of *𝒟*^*H*^ and *𝒟*^*L*^ in two subsets considering only the entries with BIT scores higher or lower than 45. Then we computed the performances in each of the four sets and we found a stronger impact of the number of effective sequences with respect to the BIT score (2.b)

We also analyzed in detail which type of contacts are better predicted. In table 1 we report the PPV for different nucleotide pairs and we can clearly see that C:G and A:U, that (mainly) correspond to canonical base pairs, are usually well predicted with a PPV of 75% and 65% respectively. These contact types are much better identified than the other contacts since the physical interaction between them is stronger and as consequence also the co-evolutionary signal. Note that the fact that C:G pair is more stable than A:T could be related to the slight difference between their prediction accuracy. There are however also non-canonical pairs that are relatively well predicted, even if to a much less extent, such as the G:U pairs with a PPV of 32%.

To asses more in deep the ability of the DCA methods to predict the more challenging non-WC long-range 3D contacts, that give important information regarding the three-dimensional structure of RNA molecules, we repeat the analysis shown above but excluding from the experimental RNA map, all contacts that are in a 5×5 windows centered at any WC base pairs.

In table (2) we report the PPV for these type of contacts at *L*/10 numbers of contacts. We can immediately observe that the values are much smaller than in the case in which all residues are considered. There is essentially no signal in the *𝒟*^*L*^ set while for *𝒟*^*H*^ the PPV is between 20% and 25% with the plmDCA method EVcouplings that reaches the best performance.

**Table 2:** Performance of the DCA-based methods analyzed on the different datasets.

| Methods | $\text{TOP}_{\mathcal{D}} L/2$ | $\text{TOP}_{\mathcal{D}^H} L/2$ | $\text{TOP}_{\mathcal{D}^L} L/2$ |
| --- | --- | --- | --- |
| pydca [35] | **43.0%** | 60.3% | 24.2% |
| EVcouplings [31] | **44.8%** | 62.3% | 25.8% |
| Boltzmann Learning[42] | **43.1%** | 61.0% | 23.7% |
| GREMLIN [44] | **41.1%** | 57.5% | 23.3% |
| CCMpred [45] | **42.2%** | 59.4% | 23.6% |
| PSICOV [46] | **41.9%** | 59.1% | 23.1% |

Finally, we test the computational efficiency of different methods by assessing their runtime for the complete set of RNA structures *𝒟*. We run all tests on a Intel i7-7700 six-core processor.

As we can see from (4), the mean-field DCA in pydca and EVcouplings are the fastest approaches with a global run-time for all structures of *𝒟* of about 10/21 minutes. They are about 5 times faster than CCMpred, a pseudolikelihood based methods known to be particularly performing when optimized on GPU-based architecture, and from 15 to 30 faster than GREMLIN and PSICOV. The slowest method is the Boltzmann Learning that is about 300 times slower than mean-field DCA. Note that, as shown in Table (4), the methods tend to have two bottlenecks in terms of run-time, the first is for long RNA sequences such as the one show that is the large ribosomal subunit from *Haloarcula maris-mortui* while the second one is for deep MSA such as the family RF00163 with more than 3 *×* 10^5^ RNA sequences.

**Table 3:** Accuracy of the different DCA-based methods for the prediction of the long-range tertiary contacts.

| Methods | $\text{TOP}_{\mathcal{D}} L/10$ | $\text{TOP}_{\mathcal{D}^H} L/10$ | $\text{TOP}_{\mathcal{D}^L} L/10$ |
| --- | --- | --- | --- |
| pydca [35] | **10.9%** | 18.8% | 2.4% |
| EVcouplings [31] | **14.9%** | 24.3% | 4.4% |
| Boltzman Learning [42] | **13.0%** | 22.4% | 2.8% |
| GREMLIN[44] | **10.8%** | 17.5% | 3.5% |
| CCMpred [45] | **11.3%** | 18.3% | 3.6% |
| PSICOV [46] | **12.0%** | 20.3% | 2.9% |

**Table 4:**
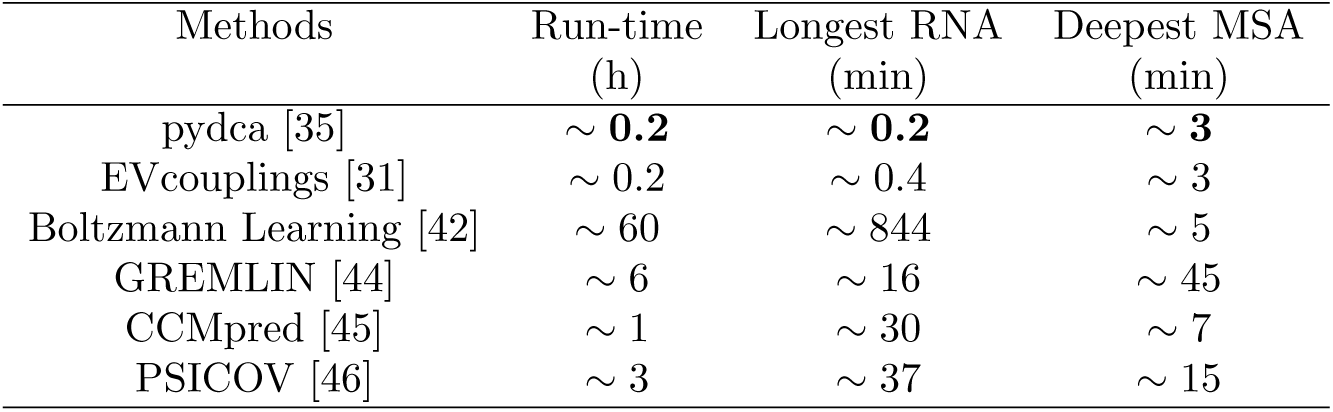
Run-time comparison of the different DCA-based methods. The longest RNA analyzed is the large ribosomal subunit from *Haloarcula marismortui* with a length of *N* =496 (RFAM RF02540) while the deepest MSA corresponds to the synthetic Hammerhead Ribozyme whose family RF00163 has more than 3 × 10^5^ RNA sequences.

### Contact type and prediction robustness

We test the robustness of the DCA-based contact predictions with respect to varying contact definitions. In table 5, we compare the PPV values of mean-field DCA for the six different contact definitions that have been introduced in Methods.

**Table 5:** Accuracy of the mean-field DCA for different contact definitions classified according to the distance threshold and the atoms used in the computation of the nucleotide pair distance.

| Contact type | N1-N9 | C1'-C1' | All | All | All | All |
| --- | --- | --- | --- | --- | --- | --- |
| Dist. Threshold (Å) | 9.5 | 12.0 | 3.5 | 5.5 | 7.5 | 9.5 |
| PPV | 43.0% | 45.9% | 41.2% | 48.1% | 54.0% | 57.0% |

Regarding the type of contacts analyzed, we see that there is no substantial difference in considering different type of contact criteria N1-N9 (9.5Å), All atoms (3.5Å) and C1’-C1’ (9.5Å) where the threshold distances have been taken from [48]. For this reason we took, as criteria along all the paper, the distance between N1 atoms for purine and N9 atoms for pyrimidine that are the atoms that established the glicosylic bond with the C1’ atoms of the pentose sugar.

Non-surprisingly, the PPV accuracy improves if the distance criteria is relaxed. For example using all atoms distance from 3.5 till 9.5 we have a PPV that increases from about 40% till a values close to 60%.

### RFAM family construction, sequence alignment and trimming

In this section we analyze how the preliminary steps of the computation, *i*.*e* the search for homologous sequences and the MSA curation, impact on the DCA prediction of RNA contacts. As a first step, since almost half of the RFAM considered do not have a large enough *M*_*eff*_ value, we modify the E-value cutoff used to constitute the RNA families in Rfam 14.1. We did it employing the cmsearch option of Infernal [34] without modifying the covariance models of the family but choosing a large E-value threshold equal to 0.99. On the other side, since the introduction of too many sequences in a given family can introduce noise, we also repeated the same analysis but with a more stringent cut-off of 0.0001.

The results do not change significantly but we can observe few trends : the enlarging of the thresholds slightly improves the contact prediction of families with *M*_*eff*_ less then 70 while keeping constant the prediction of the other ones. A more severe cut-off makes instead the performance predictions lower of about 2%.

The way in which the alignment is performed impacts the prediction performances more substantially. Alignments obtained via ClustalW lead to less accurate PPV values of about 20%. MUSCLE and MAAFT perform better than ClustalW with more or less the same accuracy (PPV values of about 30%). Finally, alignments done using Infernal improve substantially the performance with a PPV score that is about 10% above those obtained using MUSCLE and MAAFT. The higher PPV values from Infernal are, however, not surprising, as the covariance models used in Infernal are constructed from seed alignments that in turn are constructed using available information and annotations about RNA sequences such as RNA 2d structure.

Finally, the way in which the alignment is trimmed also does not change the mean-field DCA performances and usually excluding the columns in MSA that have less than 50% of nucleotides results in accurate contact predictions.

### Example of contact predictions

In order to provide an example of RNA contact prediction we analyze the aptamer domain of the Adenine Riboswitch from *Vibrio vulnificus*. Its three-dimensional structure has been deposited in the Protein Data Bank with the code 4TZX [49]. This type of ncRNA that resides in the 5 untranslated region of the *add* adenosine deaminase mRNA is one of the smallest (about 120 residue) riboswitches and it controls the translation machinery. When adenine, to which it binds, is not present, the aptamer region has a fold that prevent translation initiation. In the presence of adenine, ligand-binding allosteric effects lead to the rearrangement of the secondary structure of the aptamer region and as a consequence to the initiation of translation.

In these conditions, the structure is formed by three helices P1, P2, and P3 (see fig (4)) and three loops. In physical space, three dimensional contacts occurring between stem-loop 2 and 3 stabilize the 3D structure.

In order to predict the contacts, we start from the RFAM RF00167 (BIT score 59.4) and re-align all sequences in the family using Infernal tool. We then applied mean-field DCA implemented in pydca and the results are shown in Table 7 and in Figure 4.

**Table 6:** Impact of the MSA construction, alignment and trimming on the performances of the mean-field DCA contact prediction method.

|  | Methods | TOP<sub>D</sub> L/2 | TOP<sub>D<sup>H</sup></sub> L/2 | TOP<sub>D<sup>L</sup></sub> L/2 |
| --- | --- | --- | --- | --- |
| Search | E-value 0.0001 | 41.4% | 57.8% | 23.6% |
|  | Rfam 14.1 | 43.0% | 60.3% | 24.2% |
|  | E-value 0.99 | **43.8%** | **60.5%** | **25.6%** |
| Align | ClustalW | 22.2% | 27.5% | 16.5% |
|  | Infernal | **43.0%** | **60.3%** | **24.2%** |
|  | MUSCLE | 28.0% | 37.9% | 17.1% |
|  | MAFFT | 27.5% | 37.8% | 16.3% |
| Trim | Full<sub>refseq</sub> | 43.0% | 60.3% | 24.2% |
|  | Full<sub>gap50</sub> | 43.5% | **60.7%** | 24.7% |
|  | Full<sub>gap20</sub> | **43.6%** | 60.5% | **25.1%** |

**Table 7:** Predicted Positive Values (PPV) for different number of contact *N* and different thresholds for the Adenine Riboswitch from *Vibrio vulnificus*.

| | $N=7$ | $N=14$ | $N=35$ | $N=71$ | $N=142$ |
| --- | --- | --- | --- | --- | --- |
| PPV (cut 9.5 Å) | 1.0 | 1.0 | 0.86 | 0.55 | 0.35 |
| PPV (cut 11.5 Å) | 1.0 | 1.0 | 0.89 | 0.70 | 0.51 |

**Figure 3:**
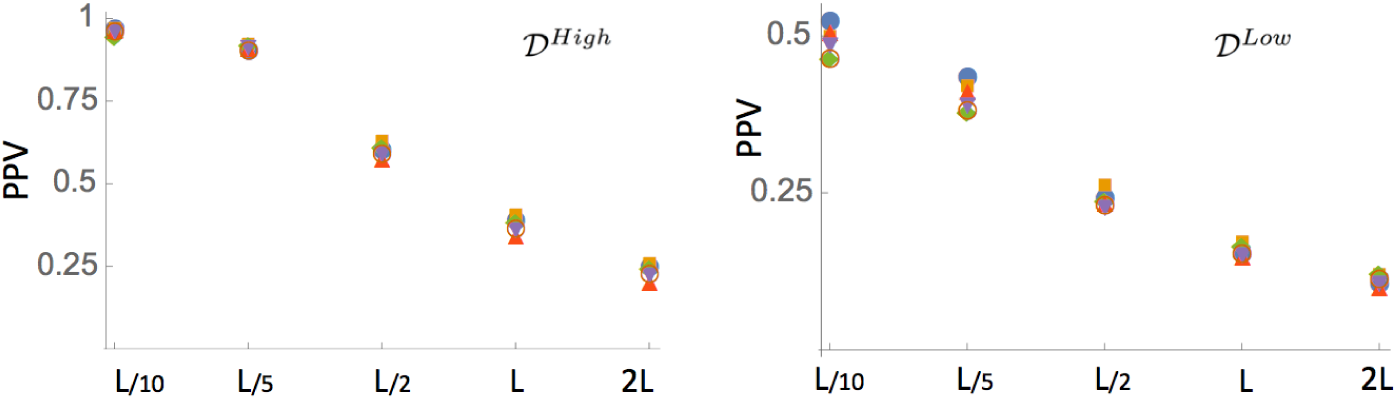
Prediction performances of the methods on the *𝒟*^*H*^ and *𝒟*^*L*^ datasets. Only contacts that are separated along the sequence of at least 4 nucleotides are considered here.

**Figure 4:**
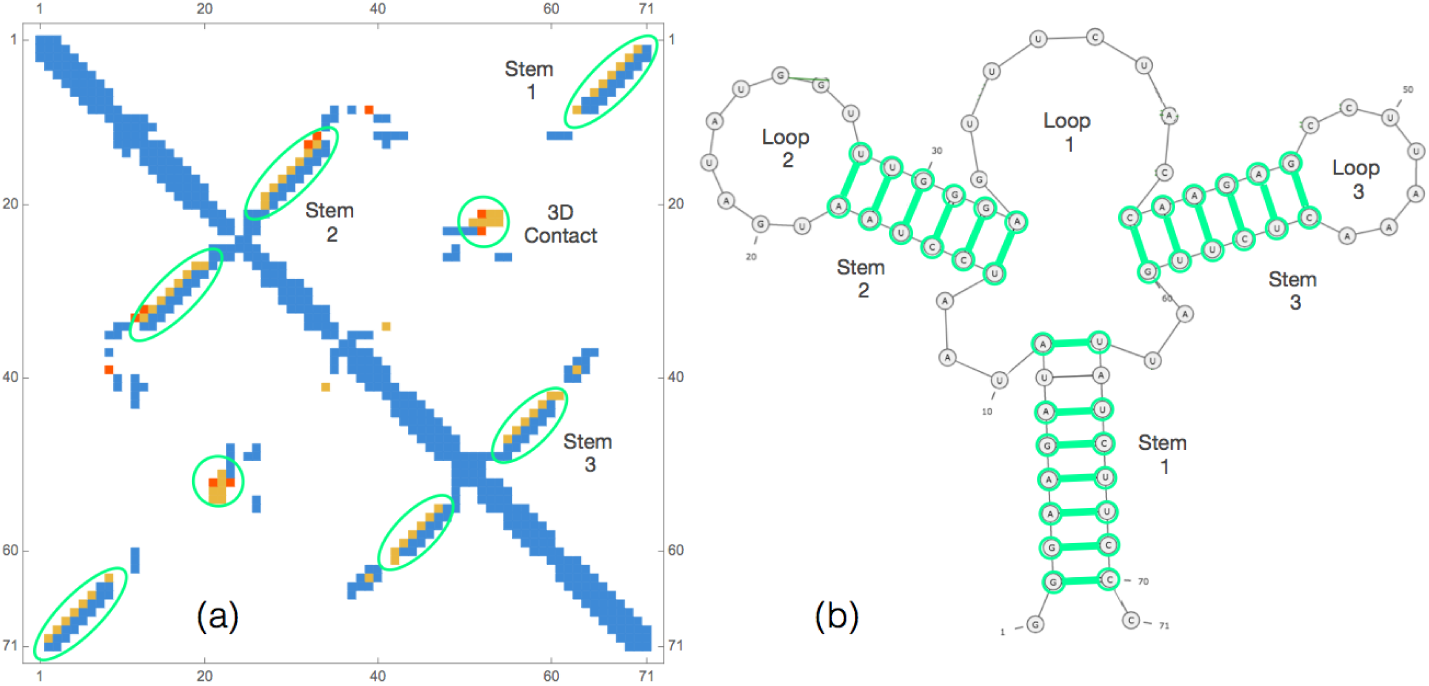
(a) Contact map of the Adenine Riboswitch from *Vibrio vulnificus* : in blue the contact from PDB struture 4TZX, in green and red the correct and wrong predicted contacts respectively (in the top 35 pairs). In (b) we plot its secondary structure with in green all correct predicted WC base-pairs in the top 35 pairs.

As we can see from 7 the PPV are quite high as all 20 but one WC base pairs of the three stems are correctly identified in the first 35 contacts (=*L*/2). Moreover there are also several 3D contacts, i.e. long range contacts in the sequence that are away from any WC base pairs, predicted. For example there are 5 contacts in the green circle of the contact map of Fig. 4 that signal an interaction between the loops 2 and 3. In total in the first 7 3D contacts, 4 of them (PPV_3*D*_ = 57%) are correctly predicted (distance threshold at 9.5 Å) but this number rises to 6 (PPV_3*D*_ = 86%) if the distance threshold is enlarged to 11.5 Å. As shown in a series of recent papers, the correct prediction of these 3D contacts and their use as constraints in molecular modeling tools can substantially improve the accuracy of the RNA 3D structure prediction [13, 26, 31, 32].

## Discussion

Co-evolution between pairs of nucleotides in MSA of homologous RNAs has shown can provide important information about the three-dimensional structure of RNA. As RNA structure and function are closely interlinked, co-evolutionary methods promise to play an important role in the understanding of a wide series of RNA-based biological mechanisms.

In order to assess the accuracy of six different widely known DCA methods more precisely, we first constructed a well curated dataset of about 70 RNA structures with good resolution. We then perform MSA alignment of their corresponding RFAM families and run six contact prediction methods : mean-field pydca, EVcouplings, Boltzman Learning, GREMLIN, PSICOV and CCMPRED.

We find that there are no statistical significant differences between their performances as measured by PPV. The prediction performance strongly depend on two factors : the first one is the number of effective sequences *M*_*eff*_ of the given RFAM family. Indeed we show that only families that have at least *M*_*eff*_ of the order of about 100 tend to lead to more reliable predictions performances. The second is the procedure used to perform the alignment. In this regard alignments done using Infernal give results much better than those obtained with other methods with the caveat that the Infernal covariance model is based on additional information.

We also noticed that the prediction of 3D contacts that are far in the sequence and from any WC base pairs, does not reach yet a satisfactory performance for the majority of the entries. While expected (these contacts should exhibit weaker signals when compared with the WC base pairs) this is also unfortunate as prediction of such long-ranged contacts can considerable boost the 3D structure prediction. Machine-learning methods could be used in this more difficult identification since these methods are constructed and optimize to detect weak signals from noisy background.

Finally, both the resolution of RNA 3D structures and the types of contact considered do not impact significantly our measure of the methods’ performance.

In summary, improving RNA contact predictions remains a challenge. The analysis done in this paper, with the construction of a new dataset of RNA structures and all tests done, provides new insights on DCA-based approaches highlighting their strong and weak points and could be a starting point for future improvements of the fields.

